# CIZ1 regulates G1 length and the CDK threshold for initiation of DNA replication to prevent DNA replication stress

**DOI:** 10.1101/2024.09.02.610838

**Authors:** James Tollitt, Tiernan Briggs, Sarah L. Allinson, Christopher J. Staples, Jason L. Parsons, Richard L. Mort, Nikki A. Copeland

**Affiliations:** Biomedical and Life Sciences, Faculty of Health and Medicine, Lancaster University, LA1 4YQ; School of Medical and Health Sciences, Bangor University, Bangor, Gwynedd, LL57 2DG, UK; Institute of Cancer and Genomic Sciences, College of Medical and Dental Sciences, University of Birmingham, Edgbaston, Birmingham B15 2TT

**Keywords:** G1 length, cyclin dependent kinase threshold, Cell cycle, DNA replication stress, CDK, Fucci(CA)

## Abstract

Eukaryotic cell division is regulated by oscillating CDK activity, which must reach critical CDK threshold activity levels to progress through cell cycle stages. In low mitogen, low CDK environments cells exit the cell cycle into a non-proliferative quiescent state, G0, that plays essential roles in stem cell maintenance and cellular homeostasis. CIZ1 regulates cell cycle and epigenetic programmes, and CIZ1 ablation enhances genomic instability after release from quiescence. Here, we determined the mechanisms that promote genome instability in CIZ1 ablated cells using a combination of Fucci(CA) live cell imaging, cell-free DNA replication assays and DNA combing. Cell cycle dynamics are unaffected in *CIZ1^−/−^* (CIZ1 KO) fibroblasts; however, a specific post-quiescent phenotype is observed resulting in a reduced G1 phase and cell cycle length. The reduction in G1 length in CIZ1 KO cells is associated with increased cyclin E1/E2 and A2 expression, and enhanced phosphorylation of Rb leading to early restriction point bypass. *CIZ1^−/−^* cells are deficient in cyclin A chromatin binding and required increased cyclin-CDK activity for the initiation of DNA replication, which is associated with DNA replication stress *in vitro* and *in vivo*. Significantly, the CDK threshold for initiation of DNA replication was 2-fold higher in CIZ1 KO nuclei than parental controls. Importantly, addition of recombinant CIZ1 *in vitro* and *in vivo* promotes recruitment of cyclin A to chromatin and reinstates the CDK threshold for initiation of DNA replication, reversing DNA replication stress and increasing replication fork rates. Loss of CIZ1 is associated with dysregulated cyclin-CDK signalling, resulting in reduced G1 length, an increased CDK activity threshold required to promote initiation of DNA replication that results in DNA replication stress. These data suggest that CIZ1 facilitates recruitment of cyclin-CDK complexes to chromatin and contributes to the mechanisms that determine the threshold CDK activity required for the G1/S transition in post-quiescent cells. Taken together the data support a role for CIZ1 in the prevention of DNA replication stress and maintenance of genome stability.

## Introduction

Eukaryotic cells use a reversible cell cycle exit, quiescence or G0, to maintain a lifelong pool of cells that can proliferate in response to mitogenic signalling to repair and maintain tissue homeostasis(1). Quiescence (G0) contributes to developmental programmes, stem cell maintenance, prevention of replicative senescence in young organisms and plays important roles in healthy aging(2). Metazoan cells exit the cell cycle and enter quiescence *in vitro* in low mitogen environments or at high cell density through contact inhibition(3). The ability of cells to exit proliferation cycles into a quiescent state (G0) also has implications for therapeutic responses in cancer, where non-proliferative populations of cells are more resistant to cancer therapies (4–6). Therefore, a more complete understanding of the regulatory processes that facilitate quiescence entry and exit is required.

Here we investigate the role of CIP1 Interacting Zinc finger protein 1 (CIZ1) in cycling and post-quiescent fibroblasts. CIZ1 has multifaceted roles, contributing to epigenetic regulation (7), maintenance of X-chromosome repression and localisation (8,9), oncogenic transcriptional programs (10,11) and cell cycle regulation (12,13). CIZ1 is a CDK sensor that promotes cell cycle progression and facilitates initiation of DNA replication (14–16). CIZ1 is a non-essential gene (8,17). However, CIZ1 null mice have defects in neurological function and increased DNA damage responses(18). CIZ1 contributes to epigenetic maintenance in primary fibroblasts (19) and increase the frequency of lymphoblastic tumour formation (8,17), potentially revealing a tumour suppressor function. CIZ1 also regulates the G1/S transition through interactions with both cyclin-CDK complexes and CDK inhibitor protein p21 (14,15,20). CIZ1 promotes the G1/S transition through its interactions with cyclin E-CDK2 and cyclin A-CDK2 through localisation of their activity to chromatin to enhance the efficiency of the initiation phase of DNA replication(14) and this interaction is regulated post-translationally through CDK mediated phosphorylation of CIZ1(15).

CIZ1 ablation revealed defects in cell cycle re-entry from quiescence that underpin cellular transformation(21), however, the precise role for CIZ1 in maintaining genome integrity in proliferation-quiescence cycles is not fully understood. Quiescence is regulated by extracellular, intercellular and intracellular signalling(2,22,23). The regulation of the proliferation or quiescence decision is mediated by extracellular mitogen levels and intrinsic CDK activity levels inherited from the mother cell. There are exit points from the cell cycle into a non-replicative quiescent state in mitosis and G1 phase of the cell cycle(24,25). Intrinsic CDK activity plays a key role in the proliferation-quiescence decision and dictates the length of G1 phase, as bifurcation of CDK activity dictates whether cells rapidly exit G1 phase or at lower CDK levels enter a prolonged G1 state (25). Alterations in mitogenic signalling are integrated into cell cycle proliferation decisions providing a memory of mitogen exposure through alteration of cyclin D levels (26). Intracellular CDK activity is regulated by the balance of CDK activity and CDK inhibitor protein levels that provide a molecular memory of genotoxic stress from previous cell cycles(25,27–30).

In mammalian cells, the restriction point (R) determines the phase of G1 at which cells no longer require mitogens to complete the cell cycle (31–34). The mechanistic basis of the restriction point involves the integration of extracellular and intracellular signalling, including transcriptional regulation via the RB-E2F pathway and intracellular CDK activity (23,35). There is a precise CDK threshold at which cells bypass R and progress through the cell cycle, and the level of intracellular CDK activity in G1 phase accurately predicts the cells that will bypass R in the absence of mitogenic signalling(36). Intracellular CDK activity controls the relative depth of quiescence, with low CDK activity cells leading to a deeper quiescent state or potentially senescence whereas higher CDK activity cells are more likely to rapidly exit quiescence. (23) (37).

Here we show that CIZ1 regulates cell cycle dynamics in post-quiescent cells. Comparison of parental and *CIZ1^−/−^* cells shows that CIZ1 contributes to mechanisms that regulate G1 length and establish the CDK threshold for G1 exit and initiation of DNA replication. The loss of CIZ1 promotes dysregulation of cyclin E expression and cyclin A localisation resulting in profound effects on restriction point timing and G1 length in post-quiescent cells, ultimately resulting in a DNA replication stress phenotype. Taken together, these data provide insights into the molecular function of CIZ1 in post-quiescent cells and provide mechanistic understanding of its role in maintenance of genomic stability by prevention of CDK mediated DNA replication stress.

## Methods

### Cell Culture

Cell culture was performed using standard aseptic techniques in a laminar flow hood. Mycoplasma testing of cell lines was performed routinely to ensure no contamination. Cell lines used were NIH 3T3 cells (ATCC CRL-1658) and NIH 3T3 derived *CIZ1*^−/−^ (hereafter referred to as CIZ1 KO), and CIZ1 KO cells plus stably integrated GFP-CIZ1 referred to as CIZ1 Add back (CIZ1-AB). Parental cells and CIZ1 KO cells were also produced containing the FucciCA construct (38). All cells were cultured in DMEM low glucose (1g/L glucose) Glutamax II plus pyruvate (Gibco), 10% (v/v) FBS (LABTECH), Penicillin, Streptomycin, glutamine (1X, Gibco) at 37°C. Cells were passaged every 2-3 days to ensure 70% confluency was not exceeded.

### G0 Cell Synchronisation

NIH 3T3 cells were synchronised in G0 by contact inhibition (14,15,39). At confluence, media was changed and cells cultured for a further 48 hours. Cells were released from G0 by passaging with a minimum 1:4 dilution. Cell cycle synchronisation was confirmed by EdU labelling and quantification of S phase entry. In addition, cell cycle kinetics were assessed using Fucci(CA) live cell imaging to provide complementary cell cycle kinetics data.

### CRISPR-Cas9

Cells were cultured to 70% confluence and 1 x 10^6^ cells were transfected using a Lonza nucleofector 2B, Kit R, transfection program U-30. CRISPR-Cas9 knockout was performed using the CD4 + gene art plasmid kit (Invitrogen). Cell line generation DNA sequencing confirmed frame shifts and nonsense mutations and loss of CIZ1 protein expression was confirmed by western blot. 2 CIZ1 KO clones were produced for each guide RNA (Ciz1 KO1 for5’ – TTGCTCCTACAGCAGTTGCAGTTTT-3’, CIZ1 KO1 rev 5’-TCGAACTGCTGTAGGAGCAACGGTG – 3’) and KO-2 (CIZ1 KO2 for 5’ – CAAGGTATGGCAGTTCCCCGGTTTT – 3’ and CIZ1 KO2 rev 5’ – CGGGGAACTGCCATACCTTGCGGTG – 3’).

### Fucci(CA) Cell Line Generation

Fucci(CA) expressing cell lines were generated using Tandem-Fucci(CA)(38) cloned into a piggyBac compatible plasmid. Integration was achieved using the piggyBac recombination system (40)

### Live Cell Imaging

Live cell imaging of Fucci(CA) cells was performed using a Zeiss LSM880 confocal microscope with an environmental chamber maintaining a constant temperature of 37 °C and providing 5% CO_2_ in air. Cells were plated on a CellVis 24 well imaging plate, and images acquired at 15 minute intervals. For cell cycle analysis, data was collected for 60 hours and analysed using the Fiji distribution of ImageJ using the Fucci_Tools.ijm plugin (https://github.com/richiemort79/fucci_tools)(41). For serum removal experiments, serum containing media was removed, cells washed in PBS and serum free media added. Images were taken from 17 hours post release until 26 hours.

### Transfection (Plasmid DNA)

Transfection was performed using the Nucleofector system (Lonza). Transfection Amaxa® Cell Line Nucleofector® Kit R was used with 1 microgram of plasmid following recommended procedures. Transfection reagents were equilibrated to room temperature for 30 minutes prior to transfections. Cells were transfected using program U-30. Cells incubated in fresh media for 48 hours and media plus selection was added for all further passages. For CIZ1 WT and CIZ1 KO cell lines plus Fucci(CA) puromycin was added at 1 μg/ml (Gibco) and CIZ1 AB cells G418 (Gibco) was added at 500 μg/ml.

### EdU labelling

Cells were grown in plates containing autoclaved coverslips and incubated with 10 µM EdU (Thermo Fisher Scientific) for 30 minutes. For synchronised cells post-quiescence, EdU was added at 15 hours post release and imaged at indicated timepoints. For Restriction point analysis, EdU was added after addition of serum free medium and EdU labelled cells quantified at 24 hours after release from quiescence. For all experiments, coverslips were removed, and washed 3 x 3 minutes in 1 ml PBS. Coverslips were fixed with 0.2 ml 4 % PFA in PBS for 15 minutes at room temperature. PFA was removed and coverslips washed 3 x 3 minutes with 1 ml PBS. EdU was fluorescently labelled using the Click-iT™ EdU Cell Proliferation Kit for Imaging (Thermo Fisher Scientific). Alexa Fluor 555 azide and Alexa Fluor 488 azide were used as indicated. After labelling and washing coverslips were mounted onto glass slides with VECTASHIELD® Antifade Mounting Media with DAPI (Vector Laboratories). Slides were imaged using a Zeiss Axiozoom fluorescence microscope and 100 nuclei counted for each experiment and all experiments performed in triplicate.

EdU/PI labelling was performed using the Flow Cytometry was performed using the Click-iT™ EdU Cell Proliferation Kit (Invitrogen). Coverslips were mounted in vectashield with DAPI (VectaLabs). For analysis of post-quiescent cells, EdU was cultured continuous with the cells after addition of fresh media.

### Immunofluorescence

Cells were cultured on coverslips. For total protein, cells were washed in CSK buffer and fixed in 4% PFA for 10 minutes. For chromatin samples, cells were permeabilised in 0.5% Triton-100 in CSK buffer (10 mM PIPES-KOH pH6.8, 100mM NaCl, 300 mM sucrose, 1mM EGTA, 1mM MgCl_2_, 1 mM DTT) prior to fixation in 4% PFA for 10 minutes. Samples were blocked in 3 % BSA-PBST. Primary antibodies were incubated overnight at 4°C. Coverslips were washed, and secondary antibody was added for 1 hour at room temperature. These steps were repeated for combing labelling with multiple rounds of primary antibody binding. Coverslips were mounted in VectaShield with DAPI (for cell immunofluorescence) or prolong anti fade gold (DNA Combing).

### Flow cytometry

Cells were incubated with 10 µM EdU for 1 hour, cells were harvested by trypsinisation. Trypsin was quenched with DMEM, cells. Centrifuged at 500 x *g* for 5 minutes. Cells were washed 3 times in 1 % BSA in PBS centrifuging for 5 minutes at 500 x *g* after each wash. Cells were permeabilised in 0.5 % Triton X-100 in PBS for 20 minutes at 4 °C. Cells were centrifuged and washed 3 times in 1 % w/v BSA PBS and centrifuged after each wash. Cells were labelled with 500 µl of EdU labelling cocktail (from Alexa Fluor 488 azide Click-iT™ EdU Cell Proliferation Kit) for 30 minutes at 4 °C. Cells were washed 3 times in 0.1 % Triton X-100 in PBS and centrifuged after each wash. Propidium Iodide was added (50 µg/ml). Cells were imaged using a Beckman Coulter CytoFLEX. Data analysis and Figure creation was performed in the CytExpert software package (Beckman Coulter).

### Immunoblotting

Cells were harvested in PBS with 1mM DTT, 1mM PMSF and phosphatase inhibitors. Chromatin fractionation was performed by addition of Triton X-100 on ice for 5 minutes followed by 14,000 x *g* centrifugation (chromatin pellet, cytosol supernatant). Supernatant removed and an equal volume of 4x SDS-PAGE loading buffer added before boiling. Chromatin samples were resuspended up to 1x with SDS loading buffer. Samples were analysed by electrophoresis on SDS-PAGE gels (8%, 10%, or 15% v/v) and transferred to a PVDF membrane by semi dry transfer. Membranes were blocked in 1% w/v BSA in TBS-Tween (0.1%) (TBS-T). Primary antibodies were incubated overnight at 4 °C (anti-cyclin A C4710, sigma Aldrich 1:1000; cyclin E HE-12 Abcam 1:500; anti-actin A1978 Sigma Aldrich, 1:2000, anti-H3 (Abcam) 1:10000), phospho-Rb (Ser807/811) (D20B12) XP® Rabbit mAb (cell signalling #8516), Rb (4H1) Mouse mAb (cell signalling #9309). Membranes were washed in 1% w/v BSA TBS-T, secondary antibodies were incubated at room temperatures for one hour (Goat anti mouse IgG-HRP (Abcam) 1:5000; Goat anti-rabbit HRP (Sigma Aldrich) 1:5000). Membranes were washed in TBS-T and imaged by chemiluminescence using SuperSignal West *Pico* PLUS Chemiluminescent Substrate (Thermo Fisher). Gels were imaged and quantified using an iBright system (InvitroGen).

### Cell free *in vitro* DNA replication assays

Cell free DNA replication reactions were performed following the procedure outlined in (14,15). For G1 cytosolic extracts and replication competent nuclei, NIH3T3 were synchronised by contact inhibition and released for 15 hours (cytosolic extract) or 17.5 hours (replication licensed nuclei) as described(14,15). For S phase cytosolic extracts HeLa cells were synchronised by a double thymidine block. Cells were incubated with 2 mM thymidine for 24 hours, media removed, and cells released into fresh media for 8 hours and the second 2 mM thymidine treatment was performed for 16 hours. Cells were released into fresh medium for 1 hour, cells incubated at 4°C in hypotonic buffer (20 mM HEPES (pH 7.8), 15 mM potassium acetate, 0.5 mM MgCl_2_, 1 mM DTT), for 5 minutes. Cells were scrape harvested and cells homogenised in a loose dounce homogeniser (7 strokes for nuclei and 25 strokes for cytosolic extract preparation). Nuclei were isolated by centrifugation at 6000 x *g* and cytosolic extracts purified by 17,000 x g for 10 minutes. Nuclei and cytosolic extracts are flash frozen in 10 µl and 30 µl aliquots in liquid nitrogen and stored in liquid nitrogen until use.

To perform cell free DNA replication assays, nuclei were mixed with cytosolic extracts supplemented with 1:50 biotin-16-dUTP (Roche), 1:50 dilution 10 mg/ml creatine phosphokinase (Calbiochem), 1:50 1 M MgCl_2_, 10x premix (400 mM HEPES, pH 7.8, 70 mM MgCl2, 10 mM DTT, 400 mM phosphocreatine (CALBIOCHEM/MERCK), 30 mM ATP, 1 mM GTP, 1 mM CTP, 1 mM UTP, 1 mM dATP, 1 mM dGTP, 1 mM dCTP) and specified recombinant protein(s). Reactions were incubated for 30 minutes at 37 °C. For microscopy analysis, nuclei were centrifuged through a 30% (v/v) sucrose solution onto poly-D-lysine coated coverslips. Samples were washed with PBS, and antibody buffer (PBS pH7.4, 0.1% v/v Triton X-100 and 0.02 %w/v SDS), labelled for 30 minutes with streptavidin Alexa Fluor 555, washed again with antibody solution, and PBS, then mounted using VectaShield with DAPI. Cell free DNA replication assays that were used for immunoblotting, cell free reactions were performed, and reactions resuspended in an equal volume of 4X SDS-PAGE loading buffer, boiled and analysed by SDS-PAGE and Western Blotting.

### Bacterial Transformation

Competent Top10 *E. coli* was transformed on ice for 30 minutes. Plasmid DNA was added and incubated on ice for 30 °C minutes. Cells were heat shocked at 42 °C for 45s and placed back on ice for 2 minutes. 500 µl of prewarmed SOC media was added and placed in a shaking incubator at 37°C for 2 hours. Outgrowth was plated on prewarmed LB agar plates supplemented with appropriate selection antibiotics.

### Whole plasmid Site directed mutagenesis

Primers used for site directed mutagenesis of GFP-CIZ1 and CIZ1-N471-GST to remove the Cy-II box(14) were GGCGAGGCAGGCACAGACACAG and AGCGCTGGCTGGGTCTGGATCTG. PCR reactions followed the QuikChange site directed mutagenesis kit (Stratagene). PCR reactions were digested with *DpnI* (NEB), and the unmethylated plasmid DNA was used to transform TOP 10 competent *E. coli* cells. Single colonies were cultured, plasmids purified, and mutation confirmed by DNA sequencing (MWG Eurofins).

### RT-qPCR

Cell samples were harvested for RNA preparation by trypsinisation, reaction quenched in media and cells collected by centrifugation. RNA samples were isolated using the PureLink™ RNA Mini Kit (Invitrogen) and stored at −80°C. RT-qPCR was performed using Taqman primers from Invitrogen: cyclin D1 (Mm00432359_m1), cyclin E1 (Mm01266311_m1), cyclin E2 (Mm00438077_m1), cyclin A2(Mm00438063_m1) and GAPDH (Mm99999915_g1). RT-qPCR was performed using the Luna® Universal Probe One-Step RT-qPCR Kit (New England Biolabs), on a BioRad C1000 CFX96 touch screen thermal cycler. ΔΔCt values were calculated and displayed as relative quantitation (RQ).

### DNA combing

Silanized coverslips were prepared using the method described in Schwob et al. (2008). DNA was isolated from 1.2*10^7 cells and purified within agarose plugs using the BIORAD CHEF genomic DNA plug set. Plugs were incubated overnight in proteinase K at 50°C and washed 4 times in TNE50 (incorporating 1mM PMSF on the third wash). The day before combing, plugs were incubated for 30 minutes in MES-EDTA pH 5.7, then 30 minutes at 65°C in MES-EDTA pH 5.7, then plugs were incubated overnight at 42°C degrees in MES-EDTA containing β-agarase. Silanized coverslips were placed in DNA solution for 10 minutes and withdrawn at a constant rate of 18 mm/min. Coverslips were incubated at 50 °C for 1 hour then prepared for imaging by immunofluorescence.

### Antibody Labelling of IdU, CldU, and ssDNA

After DNA combing onto coverslips, all wash steps were performed in Coplin jars. Coverslips were incubated at 60 °C for 1 hour. DNA was denatured in 0.5 M NaOH for 30 minutes at room temperature. Slides were washed 3 x 3 minutes in washing buffer (0.1 % v/v Tween 20, PBS, pH 7.4) slides transferred to a humidity chamber and blocked in 200 µl blocking buffer (3% BSA, 0.1 % v/v Tween 20, PBS pH7.4) at 37 °C for 1 hour. Blocking buffer was removed and 50 µl of primary antibody solution 1 was added (Table 1). Primary antibody solutions were incubated overnight at 4 °C.

**Table 1.** Antibodies used for DNA combing.

| Solution | Antibody | Species Raised | Concentration | Supplier |
| --- | --- | --- | --- | --- |
| Primary Antibody<br>Solution 1 | IdU | Mouse | 1/20 | BD Biosciences<br>347580 |
|  | CldU | Rat | 1/125 | Serotec OBT0030 |
| Primary Antibody<br>Solution 2 | ssDNA | Mouse | 1/50 | Merck Millipore<br>MAB3034 |
| Secondary Antibody<br>Solution 1 | Anti-Rat IgG<br>Alexa Fluor 488 | Chicken | 1/50 | Thermo Fisher<br>Scientific A-21470 |
|  | Anti-Chicken IgY<br>Alexa Fluor 488 | Goat | 1/50 | Thermo Fisher<br>Scientific A-11039 |
|  | Anti-Mouse IgG<br>Alexa Fluor 633 | Goat | 1/50 | Thermo Fisher<br>Scientific A-21050 |
| Secondary Antibody<br>Solution 2 | Anti-Mouse IgG<br>Alexa Fluor 568 | Rabbit | 1/50 | Thermo Fisher<br>Scientific A-11061 |
|  | Anti-Rabbit IgG<br>Alexa Fluor 568 | Goat | 1/50 | Thermo Fisher<br>Scientific A-11036 |
| Primary antibody solution 3 | Digoxigenin | Mouse | 1/50 | Abcam (AB420) |
| Secondary antibody solution 3 | Anti-Mouse IgG<br>Alexa Fluor 633 | Goat | 1/50 | Thermo Fisher Scientific A-21050 |

Slides were washed 3 x 3 minutes in washing buffer, transferred to a humidity chamber, 50 µl of secondary antibody solution 1 (Table 1) added and incubated at 37 °C for 30 minutes. Slides were washed 3 x 3 minutes in washing buffer, transferred to a humidity chamber and 50 µl of primary antibody solution 2 was incubated for 2 hours at room temperature. Slides were washed 3 x 3 minutes in washing buffer, 50 µl of secondary antibody solution 2 was added and incubated for 30 minutes at 37 °C. Slides were washed 3 x 3 minutes in washing buffer and slides mounted on a 22 x 22mm square coverslip with ProLong™ Gold Antifade Mount (Thermo Fisher Scientific).

### Statistical analysis

For analysis of G1 length and cell cycle length for WT vs CIZ1 KO cells. Data were assessed for normality using D’Agostino & Pearson and Anderson-Darling test at 95 % confidence limits and were found to be normally distributed. Data was assessed using a 2 tailed unpaired T-test using GraphPad Prism software. RT-qPCR was analysed using GraphPad Prism using a 2-way ANOVA followed by pairwise Fisher’s LSD comparison tests. DNA combing data were assessed for normality and were not found to be normally distributed (Figure 6G). Data were analysed using non-parametric ANOVA analysis using the Kruskal-Wallis’s test followed by Dunn’s multiple comparisons test using GraphPad Prism software. Where *=p<0.05, ** = p<0.01, *** =p<0.001 and **** = p<0.0001. Cell free DNA replication analyses found data were normally distributed and analysed by two-way ANOVA followed by pairwise post hoc tests using Tukey’s Honestly Significant Difference multiple comparisons. Where *** =p<0.001 and **** = p<0.0001

## Results

### CIZ1 ablation reduces G1 and cell cycle length in post-quiescent cells

*CIZ1^−/−^* cells were produced using CRISPR-Cas9 (CIZ1 KO) in murine NIH3T3 fibroblasts (Figure S1A). CIZ1 KO clones were identified by western blotting (Figure S1B) and frame shifts confirmed using nested PCR and DNA sequencing to confirm *CIZ1* knock out (Figure S1C).

In verified CIZ1 KO cells lines, the tandem Fucci(CA) bi-cistronic cell cycle biosensor(38) was stably integrated using PiggyBac integration (42) to facilitate cell cycle analyses. The Fucci(CA) system encodes truncated versions of CDT1 and Geminin fused to mCherry and mVenus and regulated by CUL4 and APC-CDH1 ubiquitin ligases respectively (Figure 1A). Degradation of CDT1 enables visualisation of the G1/S transition, as CDT1 is degraded by activated SCF-CUL4. The stable expression of mVenus-Geminin from S-phase to mitosis following inactivation of APC-CDH1 provides a second marker of the G1/S transition (Figure 1B, C). Analysis of exponentially dividing WT parental cells and CIZ1 KO cells by Fucci(CA) live cell imaging enabled precise quantification of G1 length and cell cycle length (Figure 1D; Supplementary Video 1). Interpolation of a single cell cycle, where cells are standardised by mitosis (time point 0) and followed for 1 complete cell cycle at mitosis showed similar degradation kinetics for mCherry-Cdt1 and mVenus-Geminin (Figure 1D). Quantitation of G1 length in Fucci(CA) WT parental cells and CIZ1 KO cells cycle length showed no differences of 6.8 ± 2.7 and 6.9 ±2.7 hours respectively (Figure 1E-G) or in cell cycle length (mitosis to mitosis) (Figure 1 E, F, H). Similarly, asynchronous cells showed similar cell cycle profiles by flow cytometry (Figure S2A). Collectively, the data show that CIZ1 is not essential for regulation of cell cycle length in exponentially dividing murine fibroblasts, consistent with its non-essential developmental role (8,17–19).

**Figure 1.**
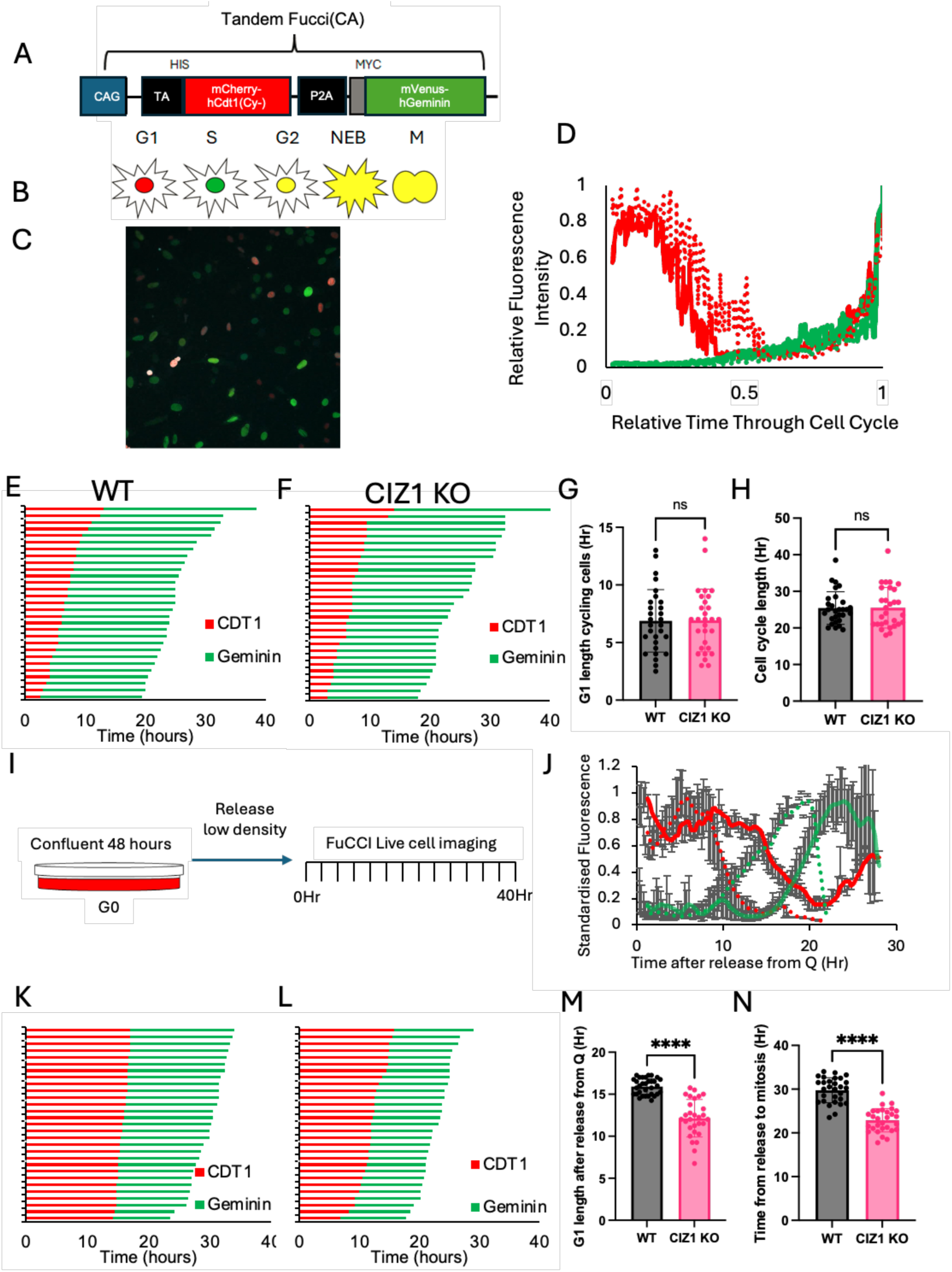
CIZ1 ablation reduces G1 length in post-quiescent fibroblasts. A) Cartoon of the bicistronic Tandem Fucci(CA) construct, showing mVenus-Geminin and mCherry hCdt1 (Cy-) fused with the porcine teschovirus-1 2A peptide sequence. B) Colour scheme for each cell cycle phase for Fucci(CA). C) Representative fluorescence microscope image of Fucci(CA) labelled murine fibroblasts. D) Interpolated fluorescence profile for WT and CIZ1 KO fibroblasts in cyclin cells. 0 hours refers to mitosis, allowing alignment of cell cycle phases. Data show mean fluorescence for mCherry-Cdt1 (red and mVenus-Geminin (Green) for WT (solid lines) and CIZ1 KO cells (dashed lines). E) Plot of WT and F) KO cells, showing mCherry-CDT1 and for mVenus-geminin fluorescence after mitosis showing G1 length (red) and S-M length (green). G) Plot of median G1 length, showing single cell median degradation of mCherry-Cdt1 and mVenus-Geminin accumulation. Bar charts show mean value and standard deviation. Statistical significance was determined using 2-tailed unpaired T-Test. n=30. H) Plot of cell cycle length, showing single cell data for WT and CIZ1 KO cells respectively from mitosis to mitosis. Bar charts show mean value and standard deviation, n=30. Statistical significance was determined using 2-tailed unpaired T-Test. I) Experimental overview of cell sychronisation in G0 by contact inhibition and post-quiescence Fucci(CA) cell cycle imaging procedure. J) Average fluorescence signal for mCherry hCdt1 and mVenus geminin after release from quiescence. WT cells (Solid line) and CIZ1 KO (dashed lines), showing G1 length (red, mCherry-Cdt1) and S-M (green, mVenus-geminin). Data points show mean fluorescence with standard deviation. K) Plot of representative cells for WT and L) KO cells, showing timepoint of median fluorescence decrease for mCherry-CDT1 and increase for mVenus-geminin. M) Plot of median G1 length from (K), dots show each individual cell cycle length and bar charts show mean G1 length with standard deviation. Statistical significance was determined using 2-tailed unpaired T-Test, (n=30, p<0.0001). N) Plot showing cell cycle length from (L) were dot show individual cell cycle length and bar charts show mean value and standard deviation. Statistical significance was determined using 2-tailed unpaired T-Test (n=30, p<0.0001).

CIZ1 regulates epigenetic maintenance in mice and CIZ1 ablation causes epigenetic instability after successive quiescence proliferation cycles that underpins cellular transformation in murine fibroblasts (21). We therefore reasoned that CIZ1 may regulate cell cycle kinetics after exit from quiescence. WT and CIZ1 KO cells expressing the Fucci(CA) were synchronised in G0 by contact inhibition and released into G1 phase allowing post-quiescence cell cycle kinetics to be observed (Figure 1I; Supplementary Video 2, 3; Figure S2C-E). Monitoring interpolated mCherry-Cdt1 and mVenus-Geminin expression levels identified that CIZ1 KO cells reduce CDT1 levels and increase mVenus-Geminin expression at earlier time points relative to WT cells, consistent with a reduction in G1 length (Figure 1J). Monitoring the level of mCherry-Cdt1 in single cells allowed determination of the time point where 50% of CDT1 is degraded, identifying that CIZ1 KO significantly reduces the mean length of G1 phase relative to WT cells (Figure 1K, L, M) (WT, 15.9 ±0.9 hours vs KO, 12.14±2.2 hours, p<0.0001 unpaired two tailed t-test; Figure 1M). CIZ1 KO cells also showed a significant reduction in cell cycle length from quiescence to mitosis (WT 29.7±2.8 hours vs 22.87±2.7 hours for CIZ1 KO, p<0.0001) (Figure 1K, L, N, Figure S3). These data demonstrate that there are significant alterations to the signalling mechanisms that control cell cycle length in post-quiescent cells in CIZ1 KO cells and imply that CIZ1 contributes to mechanisms that establish both G1 and cell cycle length in post-quiescent cells.

### Ciz1 ablation results in early restriction point bypass and reduced G1 length in post-quiescent cells

G1 length plays a key role in genome stability and cell fate decisions. G1 length is regulated by intracellular CDK activity through the relative level of CDK activity and CDKI protein levels such as p21 levels (22,25,27–29,43). Dysregulation of cyclin-CDK activity can promote early transit from G1 to S-phase(44). To determine the expression levels of p21, G1 cyclins and S-phase cyclin, RT-qPCR was performed on post-quiescent synchronised WT and CIZ1 KO cells.

The expression of p21 and cyclin D1 were unaffected (Figure 2A, B) but the E2F1-3 regulated cyclins, cyclin E1, E2 and A2 were all expressed at higher levels in CIZ1 KO cells relative to WT cells. Cyclin E1 was significantly overexpressed 20 hours after release from quiescence, cyclin E2 at 20- and 24-hours post-quiescence and cyclin A2 16 hours post-quiescence (Figure 2C-E). These data suggest that CIZ1 KO cells have increased CDK activity relative to WT cells. The restriction point is a key determinant of G1 length. The timing of restriction point can be determined by identifying the point where cells become growth factor independent for completion of the cell cycle and through monitoring activity of E2F1-3 transcriptional activity(45). To investigate if the Rb-E2F pathway is dysregulated in CIZ1 KO cells, the temporal expression of 3 Rb-E2F regulated transcripts (*CCNE1, CCNE2, CCNA2*) were assessed using qRT-PCR using post-quiescent synchronised cells where serum was removed at indicated time points (Figure 2F). This showed that cyclin E1, cyclin E2 and cyclin A2 mRNA levels increased earlier and to higher levels in CIZ1 KO cells relative to WT cells (Figure 2G-I). Cyclin E1 and E2 expression was significantly increased after removal of serum at 12 and 14 hours after release from quiescence (qPCR performed at 24 hours after release) and cyclin A2 when serum was removed at 16 hours (Figure 2I-L). Consistent with an increase in cyclin E-CDK2 activity, post-quiescent CIZ1 KO cells have increased RB phosphorylation at earlier time points and at higher levels relative to WT cells (Figure 2M). These data suggest that CDK activity is dysregulated in CIZ1 KO cells that could result in early restriction point bypass

**Figure 2.**
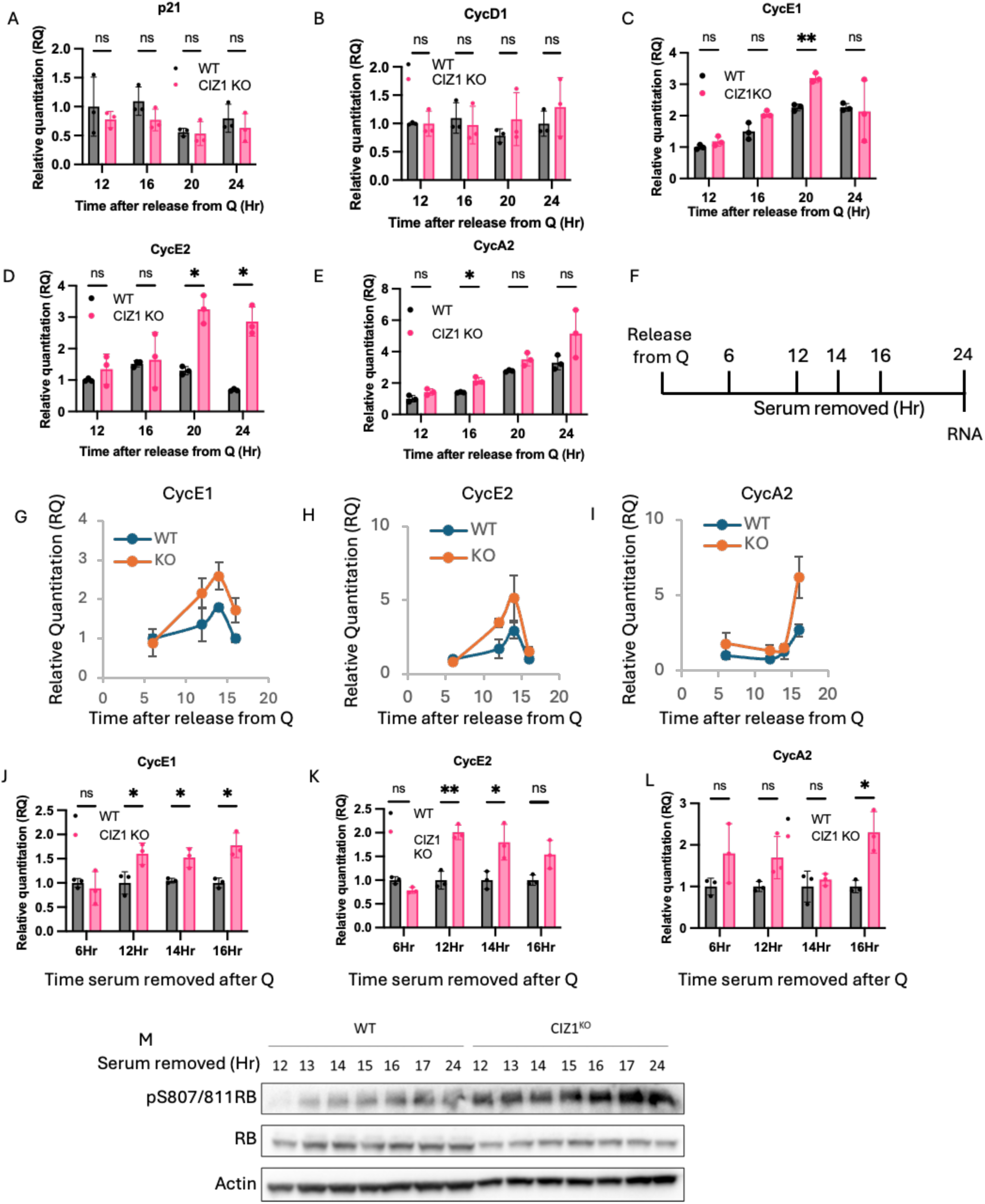
CIZ1 ablation increases cyclin E1, cyclin E2 and cyclin A2 expression in post-quiescent cells reduces G1 length through early R bypass. A-E) qPCR monitoring expression of p21 (A), cyclin D1 (B), Cyclin E1 (C), cyclin E2 (D) and cyclin A2 (E) in WT and CIZ1 KO cells in synchronous post-quiescent cells. Data show mean ± standard deviation, where n=3. Statistical analysis was performed using 2-way ANOVA Fisher’s LSD comparisons test. * - p<0.05 and ** p<0.01. F) Experimental overview. Cells were released from quiescence and serum removed at time indicated time points. mRNA levels were quantified 20 hours post release. Expression levels are show relative to actin for WT and CIZ1 KO cells. (J-L) statistical analysis of Cyclin E1 (I), cyclin E2 (J) and cyclin A2 (K) expression where serum was removed at indicated timepoints. Data show mean ± standard deviation, where n=3. Statistical analysis was performed using 2-way ANOVA Fisher’s LSD comparisons test. * - p<0.05 and ** p<0.01. M) Western blot of total cell lysates showing RB and phospho-RB levels in WT and CIZ1 KO cells. Time points indicate where serum was removed, and cell cultured in serum free media. Cells were harvested 24 hours after release from quiescence.

To determine if restriction point timing is altered in CIZ1 KO cells, WT and CIZ1 KO Fucci(CA) cells were synchronised in G0 through contact inhibition and serum deprivation, released from quiescence into complete medium. At indicated timepoints serum was removed and cells cultured in serum free medium to identify restriction point timing (Figure 3A). Cell cycle progression was monitored using live cell imaging after serum removal using live cell imaging from 17 to 36 hours post-quiescence (Figure 3B-E). Removal of serum at 11 hours blocks cell cycle progression for both WT and CIZ1 KO cells, as indicated with stable mCherry-CDT1 accumulation. However, removal of serum 13 hours post-quiescence identified a differential response in WT and CIZ1 KO cells. CIZ1 KO cells bypass restriction point at 13 hours after release from quiescence, as CIZ1 KO cells cultured in serum free medium demonstrated efficient degradation of CDT1 and accumulation of Geminin where WT cells had reduced CDT1 degradation relative to WT cells and reduced Geminin accumulation (Figure 3C).

**Figure 3.**
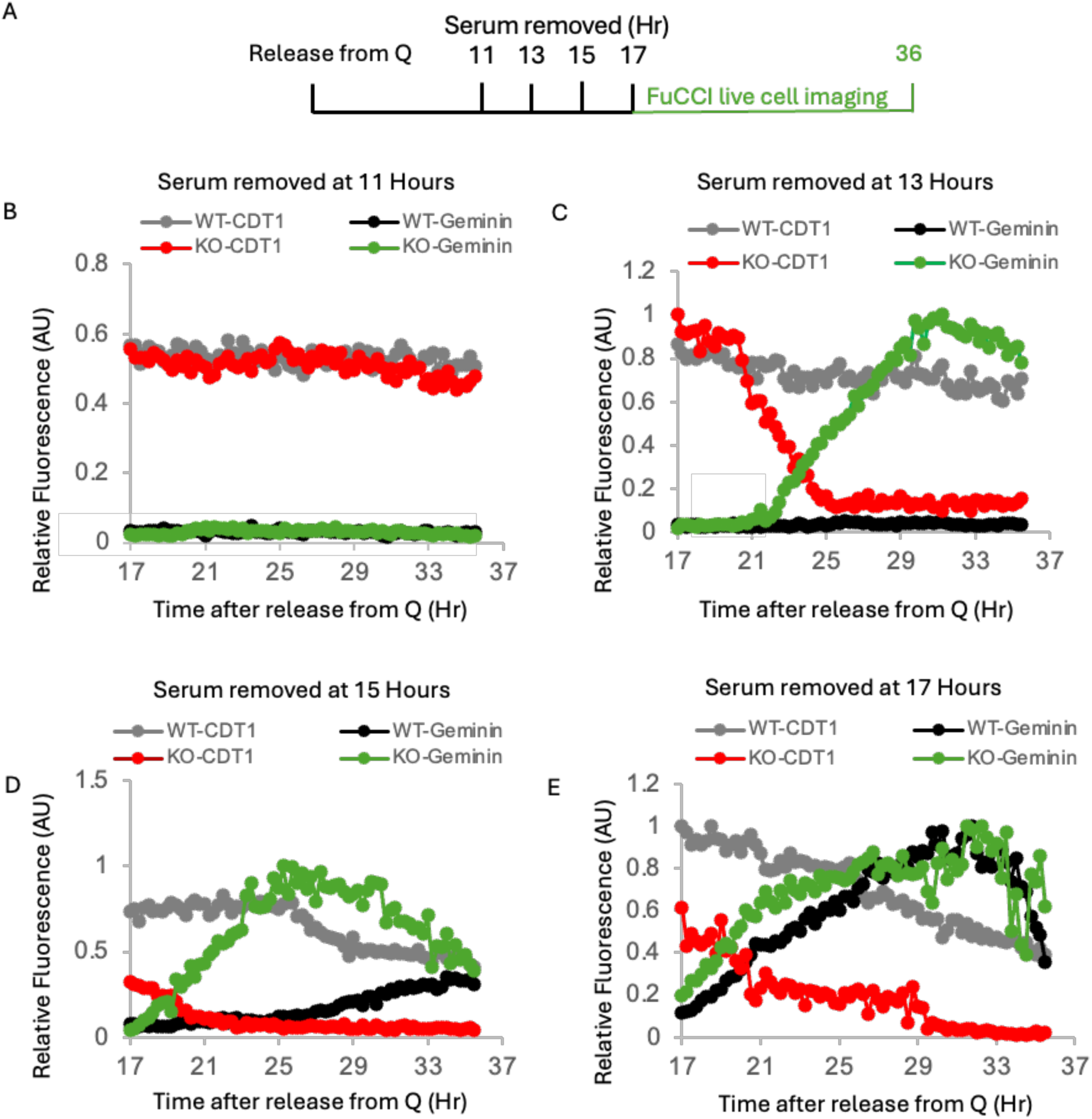
CIZ1 KO cells have reduced G1 length through earlier restriction point bypass. A) Cartoon of experimental approach. Cells were synchronised by contact inhibition and released into media plus serum. At indicated timepoints, serum was removed and live cell imaging performed from 17-36 hours. B-E) Fucci(CA) cell cycle analysis of interpolated data for WT and CIZ1 KO cells where serum removed at 11 hours (B), 13 hours (C), 15 hours (D) and 17 hours (E). mCherry-Cdt1 (grey WT, green CIZ1 KO) and mVenus-Geminin (Black WT and red CIZ1 KO) expression is shown.

Live cell imaging showed that WT cells begin to accumulate Geminin at 15 hours after release, albeit at a lower rate than CIZ1 KO cells (Figure 3D), identifying earlier restriction point timing and G1/S transition in CIZ1 KO cells. Similarly, early restriction point bypass was also evident with serum removal at 15 and 17 hours where both WT and KO cells bypass restriction point, but WT cells show a delay in Cdt1 degradation and Geminin accumulation relative to CIZ1 KO cells (Figure 3 D,E). These observations provide the molecular basis for reduced G1 length in CIZ1 KO cells, which utilise increased cyclin expression in G1 phase to bypass restriction point earlier leading to a reduction in G1 length.

### CIZ1 KO cells have reduced G1 length through dysregulation of cyclin E expression and mislocalisation of cyclin A

The observed differences in G1 length in post-quiescent WT and CIZ1 KO cells suggest that CIZ1 contributes to mechanisms that regulate G1 length in the first cell cycle following quiescence or G0. To assess if ectopic expression of CIZ1 can reverse the observed reduction in G1 length, GFP-CIZ1 was stably integrated into the chromosome of CIZ1 KO cells (CIZ1 add back, CIZ1 AB) (Figure 4A).

**Figure 4.**
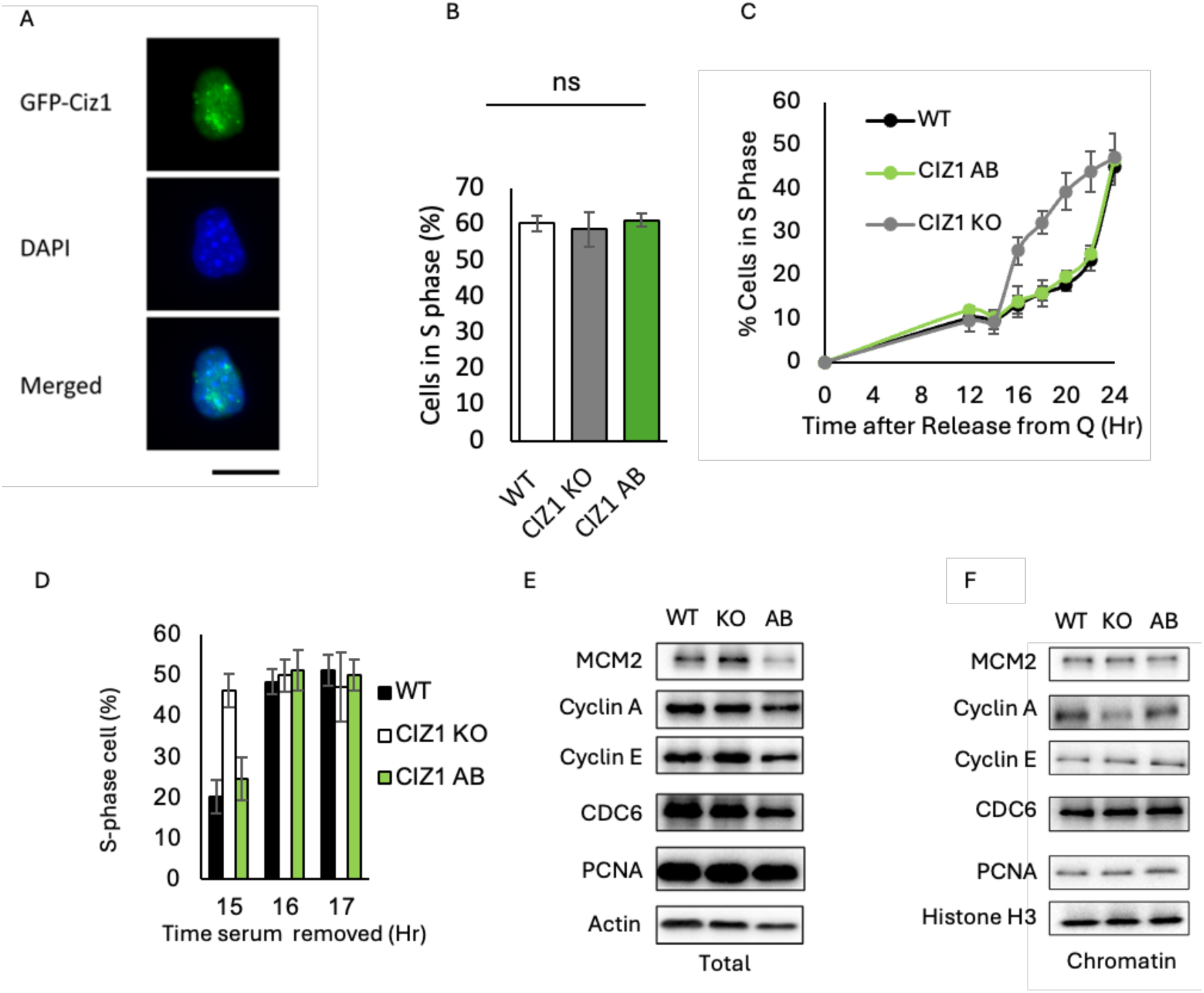
Post-quiescent CIZ1 KO fibroblasts have dysregulated cyclin dependent kinase localisation and expression. A) Fluorescence microscope of GFP-CIZ1 expressing CIZ1 KO fibroblasts, CIZ1 add back (CIZ1 AB) cells. B) Percentage of S-phase cells in EdU pulse labelled WT, CIZ1 KO and CIZ1 AB cells. C) Synchronised WT, CIZ1 KO and CIZ1 AB cells were pulse labelled with EdU, percentage of EdU labelled cells are shown from 0-24 hours after release. D) Post-quiescent cells were incubated with EdU after removal of serum at indicated time points and EdU positive cells counted at 24 hours post-quiescence. E) Western blot of WT, CIZ1 and CIZ1 AB showing cyclin A, cyclin E and actin protein levels in contact inhibited cells at day 0 (at confluence), 1 day and 2 days post confluence. 24 hours after release from quiescence is shown as a control. F) Western blot of WCE showing MCM2, cyclin A, cyclin E, Cdc6, PCNA and Actin. E) Western blot of chromatin fraction showing MCM2, cyclin A, cyclin E, Cdc6, PCNA and Actin. G) Western blot of chromatin fraction showing MCM2, cyclin A, cyclin E, Cdc6, PCNA and Actin in post-quiescent cells.

Analysis of WT, CIZ1 KO or CIZ1 AB asynchronous cells following incubation with EdU showed no significant differences in the numbers of cells in S-phase (Figure 4B). However, CIZ1 KO cells enter the S-phase several hours before WT, consistent with live cell imaging (Figure 1) and importantly, CIZ1 AB cells reversed this effect and restored G1/S transition timing in post-quiescent cells, demonstrating CIZ1 regulates G1 length and timing for S-phase entry (Figure 4C). Similarly, in post-quiescent cells the proportion of cells entering S-phase after serum removal showed that 15 hours after release CIZ1 KO cells efficiently initiate DNA replication, whereas WT and CIZ1 AB cells do not efficiently enter S-phase (Figure 4D). Next, as cyclin expression was found to be dysregulated in CIZ1 KO cells, the expression of cyclin E and A was compared in WT, CIZ1 KO and CIZ1 AB cells total protein (Figure 4E). This revealed no differences in total protein levels of cell cycle regulators or DNA replication proteins. However, CIZ1 KO cells have a reduction in cyclin A on chromatin, and this effect is effectively reversed after ectopic CIZ1 (Figure 4F). This reduction of cyclin A on chromatin is consistent with the observed role for CIZ1 in regulation of cyclin A-CDK2 localisation at the G1/S transition that is required for efficient initiation DNA replication(14).

### CIZ1 is required for efficient cyclin A chromatin localisation in CIZ1 KO cells

The length of G1 phase is controlled by intracellular cyclin-CDK activity that controls transcriptional regulation, replication licensing, and initiation of DNA at specific thresholds(12). The observed differences in cyclin E and cyclin A expression and the reduced cyclin A chromatin localisation suggests that CDK activity is dysregulated in G1 phase in CIZ1 KO cells (Figure 2 and 3). CIZ1 interacts with both cyclin E-CDK2 and cyclin A-CDK2 to promote G1/S transition(14). G1 length is a critical regulator of genomic stability. Reduction in G1 length in post-quiescent cells can under license replication origins leading to DNA replication stress (46). To establish if there is a replication licensing defect in CIZ1 KO nuclei, a cell free DNA replication assay was performed that utilises DNA replication licensed nuclei to initiate DNA replication *in vitro* (14–16,39,47). To determine if there are defects in DNA replication licensing, G1 nuclei were prepared from WT, CIZ1 KO and CIZ1 AB cells, and incubated in cytosolic S-phase extracts to promote initiation of DNA replication. Initiation of DNA replication was determined by incorporation of biotin-dUTP (Figure 5A, B). There were no observable differences in proportion of cells that initiate DNA replication for WT, CIZ1 KO and CIZ1 AB cells, suggesting that there are no gross defects in replication licensing. However, reduced cyclin A chromatin loading was found in CIZ1 KO nuclei incubated in G1 cytosolic extracts relative to WT and CIZ1 AB nuclei (Figure 5C). Cyclin A was present in the cytosolic fraction in CIZ1 KO cells, but this cyclin A pool did not efficiently load onto chromatin in CIZ1 KO cells. Incubation of CIZ1 KO nuclei with S-phase extracts showed promoted efficient chromatin loading as S-phase cytosolic extracts contain both CIZ1 and cyclin A-CDK2.

**Figure 5.**
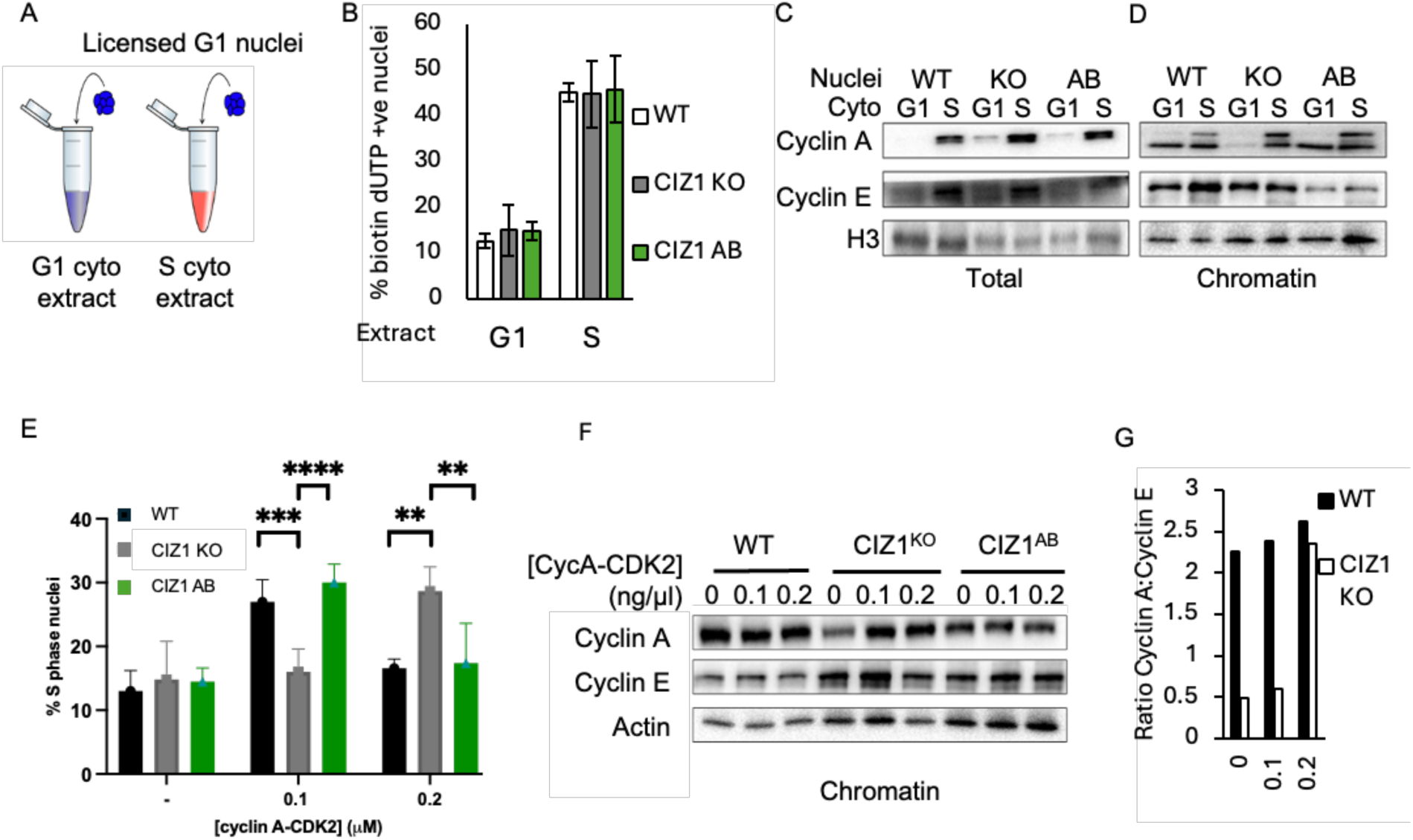
CIZ1 KO cells require increased cyclin A-CDK2 activity to promote initiation of DNA replication. A) Schematic of *in vitro* cell free reactions using licensed late G1 nuclei and S-phase cytosolic extracts. B) Initiation of DNA replication in WT, CIZ1^KO^, and CIZ1^AB^ nuclei in S phase (Biotin-16-dUTP positive) after 30-minute incubation in G1, and S phase extracts. C, D) Western blots of total protein (C), and chromatin bound fractions (D) showing cyclin E and cyclin A after incubation in G1 and S phase extracts. E) Proportion of WT, CIZ1^KO^,and CIZ1^AB^ nuclei in S phase incubation with recombinant Cyclin A-CDK2 at indicated concentrations. F) Western blot showing Cyclin A and Cyclin E in the chromatin bound fraction of WT, CIZ1 KO, and CIZ1 AB nuclei incubated with recombinant Cyclin A-CDK2 at indicated concentrations. G) Ratio of cyclin E:cyclin A on chromatin from D.

To determine how CIZ1 affects cyclin A localisation *in vitro*, cell free DNA replication reactions were performed using G1 extracts supplemented with recombinant cyclin A and CIZ1, allowing precise control of their concentration. WT, CIZ1 KO and CIZ1 AB nuclei were incubated with recombinant cyclin A-CDK2 *in vitro* at optimal (0.1 ng/μL) and high levels that prevent initiation of DNA replication (0.2 ng/μL) (Figure 5D, E). WT nuclei initiate efficiently at 0.1 ng/μL, whereas CIZ1 KO nuclei show no increase in initiation of DNA replication at this level of cyclin A-CDK2 activity. The ectopic expression of CIZ1 (CIZ1 AB) effectively compensates for ablation of CIZ1, facilitating efficient initiation of DNA replication at 0.1 ng/μL cyclin A-CDK2. Surprisingly, CIZ1 KO cells efficiently initiate DNA replication at 0.2 ng/μL cyclin A-CDK2, a level that inhibits initiation of DNA replication in WT and CIZ1 AB nuclei, identifying a shift in the CDK threshold that initiates DNA replication *in vitro*. This effect was associated with reduced cyclin A on chromatin relative to WT and CIZ1 AB nuclei (Figure 5E). At the G1/S transition, cyclin E is degraded as the S-phase E3 ligase Skp-Cullin-F box (SCF) is activated. Therefore, we determined the ratio of cyclin E and cyclin A in cell free DNA reactions. This found high cyclin E levels and low cyclin A levels in CIZ1 KO nuclei and that addition of levels of recombinant cyclin A-CDK2 that promote initiation of DNA replication restored the level cyclin A:cyclin E ratio in nuclei that efficiently initiate DNA replication (Figure 5F, G). These data suggest that cyclin A localisation is deficient in CIZ1 KO nuclei and that CIZ1 contributes to mechanisms that establish the kinase threshold to promote G1 exit and initiation of DNA replication in S phase.

### CIZ1:cyclin A stoichiometry modulates CDK threshold at the G1/S transition

To precisely determine the kinase thresholds that promote initiation, titration of recombinant cyclin A-CDK2 was performed in cell free DNA replication assays using WT, CIZ1 KO and CIZ1 AB nuclei (Figure 6A). Cyclin A-CDK2 promotes initiation of DNA *in vitro* at precise CDK activities and shows distinct concentration dependent effects with initiation of DNA replication occurring within a narrow ~2-fold range of cyclin A-CDK2 activity (Figure 6B).

**Figure 6.**
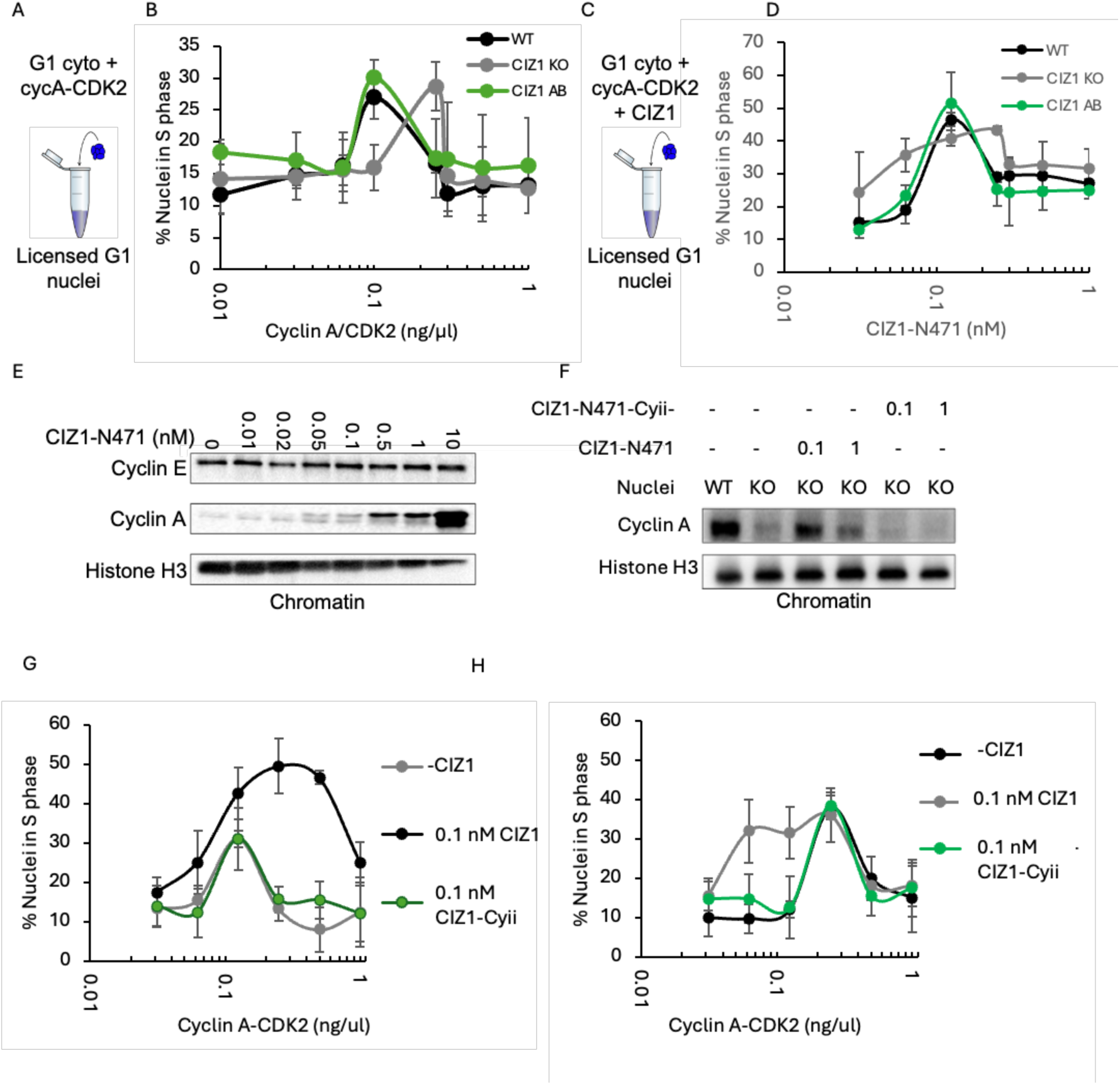
CIZ1 is required to optimise cyclin A-CDK2 activity to promote initiation of DNA replication. A) Cartoon of cell free DNA replication assays using late G1 nuclei, G1 extracts and recombinant cyclin A-CDK2. B) Effect of increasing cyclin A-CDK2 activity in WT, CIZ1 KO and CIZ1 AB nuclei. C) Cell free DNA replication assays using increasing concentrations of recombinant CIZ1 and 0.1 ng/μL. D) Effect of increasing CIZ1 on initiation of DNA replication in WT, CIZ1 KO and CIZ1 AB nuclei. E) Increasing CIZ1 titration of recombinant CIZ1 in cell free DNA replication assays promotes cyclin A binding to chromatin. F) Cell free DNA replication assays were performed with WT and CIZ1 KO nuclei and indicated WT CIZ1 or CIZ1 Cy-ii that is deficient in cyclin binding. Western blots show chromatin loading of cyclin A. G) G) Effect of increasing CIZ1 levels in WT nuclei with recombinant CIZ1 or cyclin binding deficient mutant (CIZ1 Cy-ii). H) As for G using CIZ1 KO nuclei.

In WT nuclei cyclin A-CDK2 promotes initiation of DNA replication at 0.1 ng/μL and 2-fold higher levels prevent initiation of DNA replication, returning to baseline levels. In contrast CIZ1 KO nuclei require 2-fold higher concentrations of cyclin A-CDK2 to efficiently initiate DNA replication to WT levels (Figure 6B). This level of CDK activity is non-permissive for DNA replication in WT nuclei and identifies that CIZ1 KO nuclei have an increased CDK activity threshold for passage through the G1/S transition. These results demonstrate that CIZ1 KO nuclei are replication licensed but require increased cyclin A-CDK2 to initiate efficiently DNA replication, suggesting that CIZ1 modulates the CDK thresholds required for initiation of DNA replication and the G1/S transition.

CIZ1 promotes recruitment of cyclin A-CDK2 to chromatin (14,15) and titration of CIZ1 into cell free DNA replication assays efficiently increases cyclin A chromatin binding in a concentration dependent manner (Figure 6E). To investigate if cyclin binding is required to modulate the CDK threshold levels that promote initiation of DNA replication, WT CIZ1 or a cyclin binding deficient CIZ1 mutant (CIZ1 Cy-ii)(14) were titrated into cell-free DNA replication reactions and cyclin A localisation to chromatin assessed (Figure 6F). WT nuclei have high levels of cyclin A on chromatin, whereas CIZ1 KO nuclei have very low cyclin A chromatin binding. CIZ1 enhances cyclin A chromatin loading in CIZ1 KO nuclei and this effect is dependent on cyclin binding as this effect is absent with the cyclin binding deficient CIZ1 Cy-ii mutant. To identify if cyclin binding is a key regulator to promote initiation of DNA replication, WT nuclei were incubated with increasing cyclinA-CDK2 activity with or without CIZ1 or CIZ1 Cy-ii. CIZ1 increases the permissive range of cyclin A-CDK2 activity that promotes DNA replication, suggesting that it modulates CDK thresholds at the G1/S transition and this effect is dependent on cyclin binding as this effect is absent in the cyclin binding CIZ1 mutant (Figure 6G). Titration of cyclin A-CDK2 in CIZ1 KO nuclei showed that a 2-fold higher concentration of cyclin A-CDK2 (0.2 ng/μl) is required to initiate DNA replication. The requirement for a higher cyclin A-CDK2 level is reversed by addition of recombinant CIZ1, but not by the cyclin binding deficient mutant, demonstrating that binding and localisation of cyclin A-CDK2 is required for optimal initiation of DNA replication (Figure 6H). Overall, these results show that CIZ1 KO are deficient in cyclin A chromatin loading. Importantly, the reduced levels of chromatin associated cyclin A can be reversed by ectopic expression of CIZ1, or addition of recombinant CIZ1 protein. This activity is dependent on the presence of the CIZ1 cyclin binding domain, suggesting that CIZ1:cyclin A interactions and their stoichiometry are important regulators of the CDK threshold level that promotes the G1/S transition.

### CIZ1 prevents DNA replication stress to maintain genomic stability

CIZ1 ablation in murine models increases sensitivity to genotoxic stress (18), promotes cellular transformation (21) and tumorigenesis (8,17). Here, CIZ1 KO cells have been shown to have dysregulated CDK activity through overexpression of cyclin E, mislocalisation of cyclin A with reduced chromatin binding and require increased cyclin A-CDK activity to promote initiation of DNA replication. Dysregulation of CDK activity is associated with enhanced DNA replication stress (48,49). To assess if CIZ1 KO cells have increased DNA replication stress, DNA combing was used to determine replication fork rates and origin usage (Figure 7A, B).

**Figure 7.**
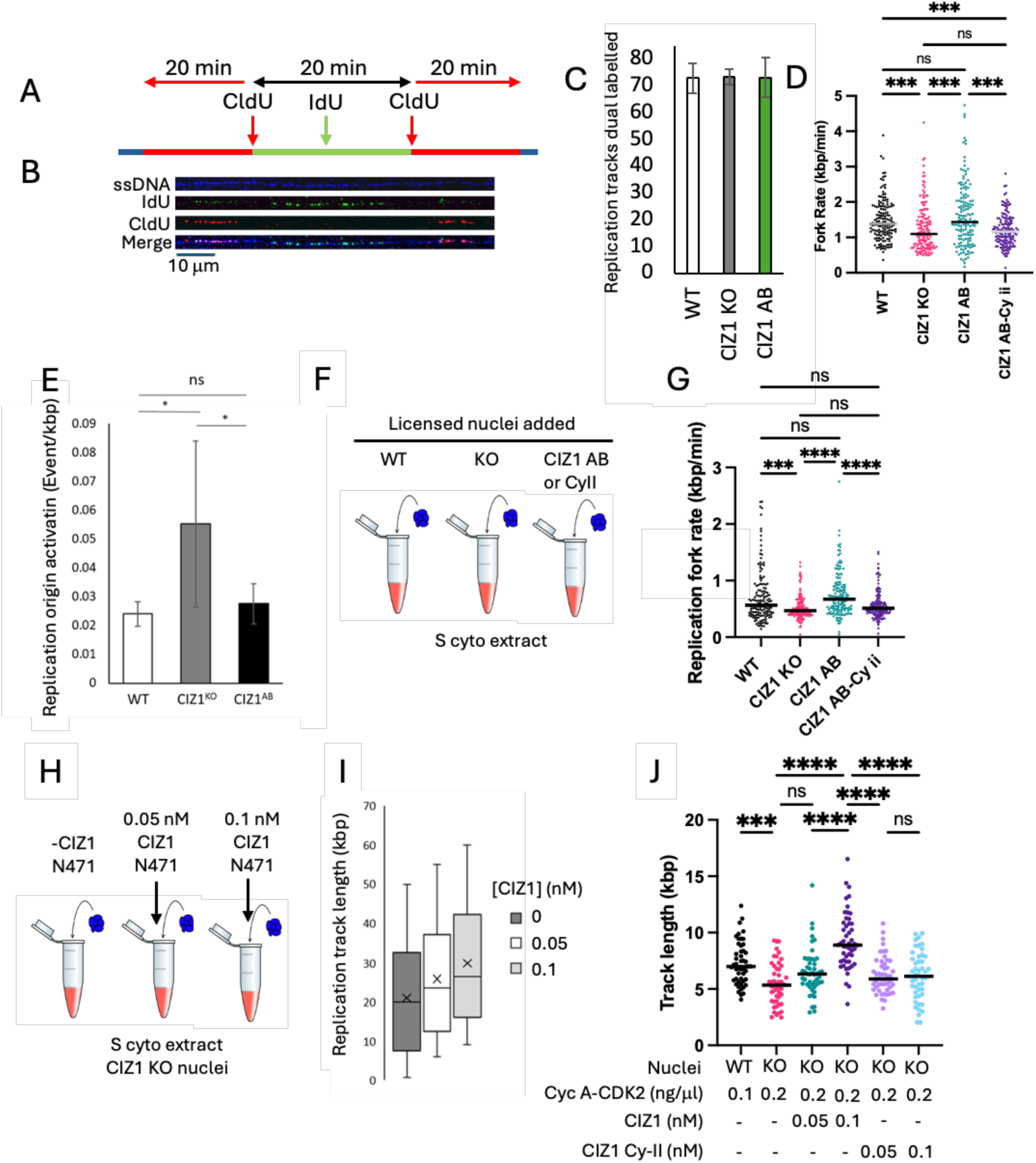
CIZ1 ablation promotes DNA replication stress. A) Experimental overview of DNA labelling procedure for DNA combing. B) Representative combed ssDNA, CldU and IdU labelled DNA. C) The percentage of WT, CIZ1 KO, and CIZ1 AB DNA fibres dual labelled after 2 consecutive 20-minute incubations in IdU and CldU. D) DNA replication fork rates were determined for WT, CIZ KO and CIZ1 AB cells. Data show individual fork rates, where n = 150. Nonparametric data was analysed by ANOVA using the Kruskal-Wallis’s test and Dunn’s multiple comparisons test in GraphPad Prism 10, where *=p<0.05, ** = p<0.01, *** =p<0.001 and **** = p<0.0001. E) DNA replication origin firing was determined per kbp of DNA. Data show mean DNA length per replication origin, with standard deviation, where n = 150 for all cells. Nonparametric data was analysed by ANOVA using the Kruskal-Wallis’s test and Dunn’s multiple comparisons test in GraphPad Prism 10. Where *=p<0.05. F) Cartoon of *in vitro* DNA replication assays using WT, CIZ1 KO and CIZ1 AB nuclei respectively in S-phase cytosolic extracts. G) Data show individual fork rates for each condition, where n = 151 for WT, 151 for KO, 154 for CIZ1 AB and 151 for CIZ1 Cy-ii. Statistical analysis was performed using non-parametric ANOVA analysis using the Kruskal-Wallis’s test and Dunn’s multiple comparisons test using GraphPad Prism software. Where *=p<0.05, ** = p<0.01, *** =p<0.001 and **** = p<0.0001. H) Cartoon of *in vitro* DNA replication assays using licensed G1 nuclei prepared from CIZ1 KO cells incubated in S-phase extracts supplemented increasing concentrations of CIZ1-N471 as indicated. I) The length of replication tracks of WT and CIZ1 KO nuclei supplemented with increasing concentrations of CIZ1-N471 or CIZ1-N471-CyII-as indicated. Data shown in box whisker plots showing median, quartiles and standard deviation. J) DNA replication track length for WT and CIZ1 KO nuclei incubated for 30 minutes in G1 extracts with recombinant cyclin A-CDK2 and CIZ1 or CIZ1 Cy-ii mutation as indicated. Data show individual fork rates, where n=50. Statistical analysis was performed using parametric 2-way ANOVA analysis using the Tukey’s multiple comparisons test using GraphPad Prism software. Where *** =p<0.001 and **** = p<0.0001.

There were no differences in bidirectional DNA synthesis (Figure 7C). However, the replication fork rate is significantly reduced in CIZ1 KO cell lines relative to the parental control cells (WT: 1.3 kbp/min, CIZ1 KO: 0.9kbp/min, p<0.001 Figure 7C). This effect is due to loss of CIZ1 as ectopic expression of GFP-CIZ1 reversed this effect (CIZ1 AB: 1.4 kbp/min, p<0.001. Figure 7D), demonstrating that CIZ1 regulates DNA replication fork rates. The reversal of DNA replication stress is dependent on cyclin binding, as the cyclin binding deficient CIZ1 mutant does not increase fork rate to WT levels as seen for CIZ1 AB cells (Figure 7D). Importantly, the loss of cyclin binding (CIZ1 Cy-ii) results in no significant differences between fork rates for CIZ1 KO and CIZ1 Cy-ii, suggesting that CIZ1 modulates fork rates through cyclin binding and chromatin localisation. To compensate for reduced fork rates, CIZ1 KO cells utilise cryptic origins increasing the frequency of replication origin firing to compensate for a reduction in fork speed in CIZ1 KO cells relative to WT and CIZ1 AB cells (Figure 7E). These data demonstrate that CIZ1 KO cells display DNA replication stress that can be reversed by ectopic expression of CIZ1. Significantly, this reversal of the DNA replication stress phenotype requires cyclin binding to CIZ1 via its Cy-ii cyclin binding box.

To complement cell-based analysis, cell-free DNA replication assays were performed and replication dynamics analysed by DNA combing. This approach allows precise titration of CIZ1 or cyclin A-CDK2 and the effect on DNA replication determined. DNA combing assays were adapted for analysis for *in vitro* DNA replication assays (Figure 7F). Critically, DNA combing from cell free experiments phenocopies cell-based experiments. CIZ1 KO nuclei produced shorter replication tracks than WT, or CIZ1 AB nuclei (Figure 7G). Furthermore, in CIZ1 KO nuclei replication tracks could be lengthened by increasing CIZ1-N471 levels in cell free assays (Figure 7 H,I) indicating that addition of recombinant CIZ1 protein could alleviate the DNA replication stress phenotype. To determine if this effect was mediated by recruitment of cyclin A-CDK2, CIZ1 Cy-ii mutant was titrated into reactions. The cyclin binding deficient mutant is unable to enhance fork rates demonstrating that cyclin binding is required to increase DNA replication fork rates and reduce DNA replication stress (Figure 7J). These data collectively demonstrate that CIZ1 KO cells have increased DNA replication stress and that this effect could be reversed through titration of recombinant CIZ1 protein or through ectopic CIZ1 expression. In both contexts cyclin binding is required to alleviate DNA replication stress, demonstrating that CIZ1 reduces DNA replication stress through binding and localisation of cyclin A-CDK2. These data suggest that CIZ1 through interactions with cyclin A-CDK2 regulates DNA replication fork speeds and contributes to mechanisms that reduce DNA replication stress to maintain genomic stability.

## Discussion

The cell cycle is governed by an increasing gradient of CDK activity. This increasing gradient of CDK activity facilitates separation of replication licensing and initiation of DNA replication, and CDK activity peaks to promote mitosis (50,51). Intracellular CDK activity is regulated at numerous levels through transcriptional regulation of cyclin expression(28), phosphatase activity(52), regulatory kinase activity (CAK/WEE1), levels of CDK inhibitor proteins (29) and ubiquitin-proteasome activity(53). These activities ensure that CDK activity is tightly regulated and coordinated within each phase of the cell cycle. Intracellular CDK activity reporters have identified that there are intrinsic CDK activity levels that dictate exit from the cell cycle or the rate of G1 exit due to stochastic CDK activities or through molecular memory by high CDK inhibitor levels in response to genetic stress in previous cell cycle. This work has identified that CIZ1 contributes to mechanisms that control the G1/S transition and modulate the specific CDK activity required for the G1/S transition and efficient initiation of DNA replication.

Analysis of CIZ1 KO fibroblasts revealed that CIZ1 modulates G1 length and affects CDK activity thresholds for the initiation of DNA replication. CIZ1 KO fibroblasts have reduced G1 length (Figure 1) and bypass restriction point significantly earlier than WT cells (Figure 2 and 3). This is mediated by increased cyclin E and cyclin A expression at earlier stages of G1 phase (Figure 2). In addition, CIZ1 KO cells also display reduced cyclin A chromatin binding (Figure 4 and 5) and increased requirement for cyclin A-CDK2 activity to promote initiation of DNA replication (Figure 6). Dysregulation of CDK activity is known to promote DNA replication stress and this was also observed in CIZ1 KO cells, demonstrating that CIZ1 contributes to mechanisms that prevent genomic instability and DNA replication stress (Figure 7). These observations provide insight into CIZ1 function and its role in genome maintenance. These data also provide insight into the association between CIZ1 ablation and the enhanced genome instability, which may underpin CIZ1 tumour suppressor function (17).

Replication licensing begins as cells exit mitosis and APC-CDH1 is activated reducing intracellular CDK activity. The length of G1 phase is an important regulatory step, as reductions in length can prevent efficient licensing of the genome (46). Here, G1 length was similar for cycling cells in both WT and CIZ1 KO cells (6.8 and 6.9 hours respectively). However, in post-quiescent cells there is a marked difference in G1 length (15.9 hours and 12.1 hours respectively) that may affect DNA replication licensing. (46). The licensing of cells can be determined *in vitro* using G1 nuclei and S-phase extracts to initiate DNA replication. The differential G1 phase length did not affect efficiency of initiation suggesting no significant defects in replication licensing.

CIZ1 KO cells show an increase in cyclin E and cyclin A expression and an increase in RB phosphorylation consistent with increased CDK activity in post-quiescent cells. The increased replication stress in CIZ1 KO cells is likely associated with the earlier overexpression of cyclin E, driving a truncated G1, resulting in DNA replication stress. Cyclin E overexpression exhausts nucleotide levels, promoting under licensing, DNA replication stress, shortening of G1, sensitivity to checkpoint inhibitors and whole genome duplication (49,54–56). Here increased cyclin E and cyclin A expression was associated with DNA replication stress, with reduced fork rates and a differential CDK threshold for initiation of DNA replication.

The differential cyclin A-CDK2 activity required for initiation of DNA replication correlates with CIZ1 KO cells have reduced chromatin associated cyclin A. The reduction in cyclin A loading to chromatin can be reversed by ectopic CIZ1 expression *in vivo* or addition of recombinant CIZ1 *in vitro* and requires the cyclin binding domain of CIZ1. In CIZ1 KO cells, DNA replication stress can be alleviated by ectopic expression of CIZ1 or addition of recombinant CIZ1 that enhances the DNA replication rate. The observed fork rate can be enhanced in vivo and *in vitro* through expression or addition of recombinant CIZ1 and significantly titration of CIZ1 levels *in vitro* led to a concentration dependent increase in fork rate. In sum, the data demonstrate that the interaction of CIZ1:cyclin A-CDK2 in G1 phase and at the G1/S transition is protective from DNA replication stress. We propose that CIZ1 modulates the CDK threshold that promotes efficient initiation of DNA replication and contributes to mechanisms that prevent DNA replication stress, thereby maintaining genomic stability.

The quantitative model for cell cycle regulation states that increasing CDK gradients are sufficient to ensure temporal control of the cell cycle. The best evidence available uses fluorescence CDK reporters to determine intracellular CDK activities through the cell cycle (27,29). They are sensitive to changes in CDK activity and to CDK inhibitor protein levels. However, they do not provide a quantitative readout for cyclin-CDK activity or levels. Here, the precise levels of cyclin A-CDK2 required to promote initiation of DNA replication was determined using an in vitro replication assay. This allows for precise biochemical analysis of the roles of CIZ1 and cyclin A-CDK2 activity (Figure 8). These data led to the following model, where CIZ1 is a key factor that determines the threshold for initiation of DNA replication and the transition from G1 to S phase. In the absence of CIZ1, G1 length is reduced in post-quiescent cells and interestingly, the reduction in G1 length is associated with an increased threshold for initiation of DNA replication relative to WT cells. This dysregulation of CDK activity promotes DNA replication stress in CIZ1 KO cells and promotes a DRS phenotype that reduces genome stability as seen elsewhere (17–19,21).

**Figure 8.**
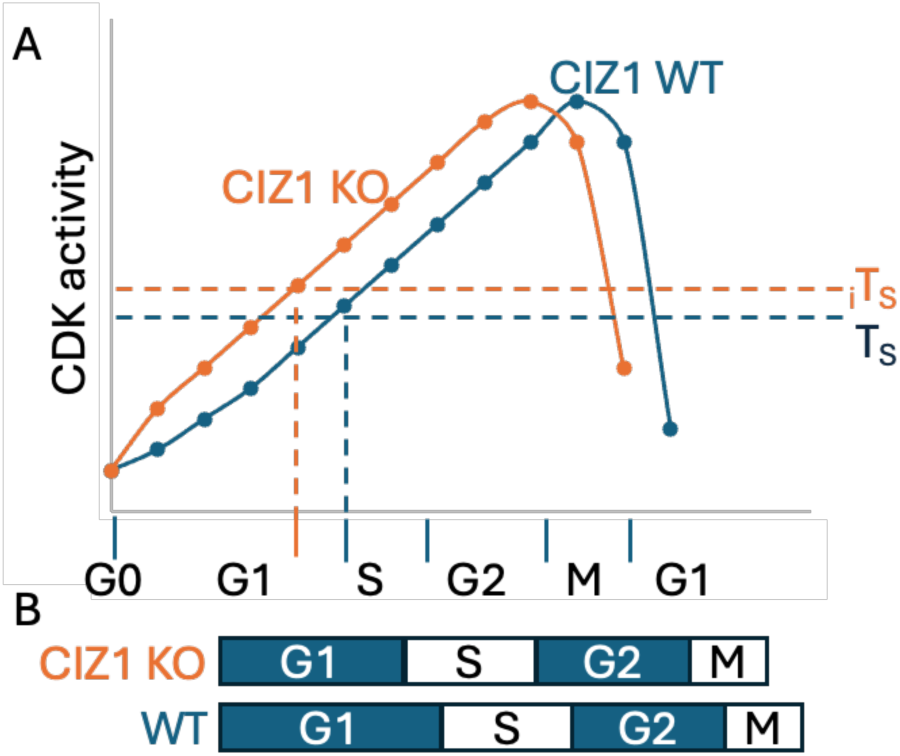
Model. CIZ1 modulates CDK thresholds to maintain genome stability. A). The quantitative model for CDK regulation of the cell cycle suggests that specific CDK thresholds regulate cell cycle transitions. The threshold for S phase entry and initiation of DNA replication is denoted T_S_ for WT cells. In CIZ1 KO cells, increased cyclin E and cyclin A expression results in increased CDK activity and a higher CDK activity threshold for initiation of DNA replication (_i_T_s_). In the absence of CIZ1, cells require a higher CDK activity to reach the threshold exit G1 phase. This is achieved through increasing CDK activity, resulting in reduced G1 length. This dysregulation of the CDK activity and mislocalisation of cyclin A in G1 phase promotes DNA replication stress in CIZ1 KO mutants. B) Differential cell cycle length in CIZ1 KO and WT cells. The increase in CDK activity in CIZ1 cells reduces G1 length and overall cell cycle length relative to WT cells, primarily through reduction in G1 length.

CIZ1 has roles in both cell cycle regulation and epigenetic maintenance. CIZ1 ablation results in alterations to the epigenetic landscape and increases propensity to generate genetic damage (9,13,19,21,57–59). At present there are 2 distinct roles for CIZ1 in epigenetic regulation. CIZ1 interacts with the lncRNA XIST that contributes to mechanisms that mediate X-chromosome inactivation and the localisation of the inactive X-chromosome (Xi) (8,9,19,57,58). In addition, it has been noted that CIZ1 also contributes to the level of polycomb repressor complex 1 and 2 (PRC1/2) mediated H2AK119Ub and H3K27me3(19). In addition, CIZ1 KO cells show defects in chromosome condensation on entry into quiescence due to reduced H4K20me1. This led to an increase in expression of homology directed repair (HDR) genes and an increase in activated Chk1 kinase and gammaH2AX levels in CIZ1 null mice (21). The results here complement and extend the mechanistic understanding of the observed genome instability in CIZ1 ablated mice. There is intrinsic DNA replication stress in CIZ1 KO cells, and the reduction in replication fork rate seen *in vivo* and *in vitro* suggest that CIZ1 plays an important role in replication fork progression or in the prevention or alleviation of DNA replication stress. The increased fork rate observed in CIZ1 KO cells when CIZ1 is ectopically expressed could indicate that global epigenetic remodelling has occurred, alleviating DNA replication stress. However, in *in vitro* DNA replication assays addition of CIZ1 protein results in increased fork rates in a concentration dependent manner, suggesting that CIZ1 is exerting its affect acutely. A second observation suggests that CIZ1 affects fork rate specifically through recruitment and localisation of cyclin A-CDK2, as fork rates are not affected by CIZ1 Cy-ii mutant that cannot interact with or recruit cyclin A-CDK2. Similarly, the requirement for cyclin A-CDK2 binding and recruitment of cyclin A to reverse this effect is demonstrated *in vivo* where ectopically expression of CIZ1 Cy-ii also does not alleviate DNA replication stress. These data do not exclude a role for the epigenetic landscape in regulation of DNA replication fork progression in CIZ1 null cells, but rather suggest that CIZ1 mediated cyclin A-CDK2 localisation is a key regulator of DNA replication fork progression. Taken together, these data suggest that CIZ1 maintains genomic stability through regulation of the CDK threshold that promotes initiation of DNA replication and regulates the rate of the DNA replication fork progression.

