## Supplementary figures and images for "CIZ1 regulates G1 length and the CDK threshold for initiation of DNA replication to prevent DNA replication stress"

### Figure S1

#### Slide 1
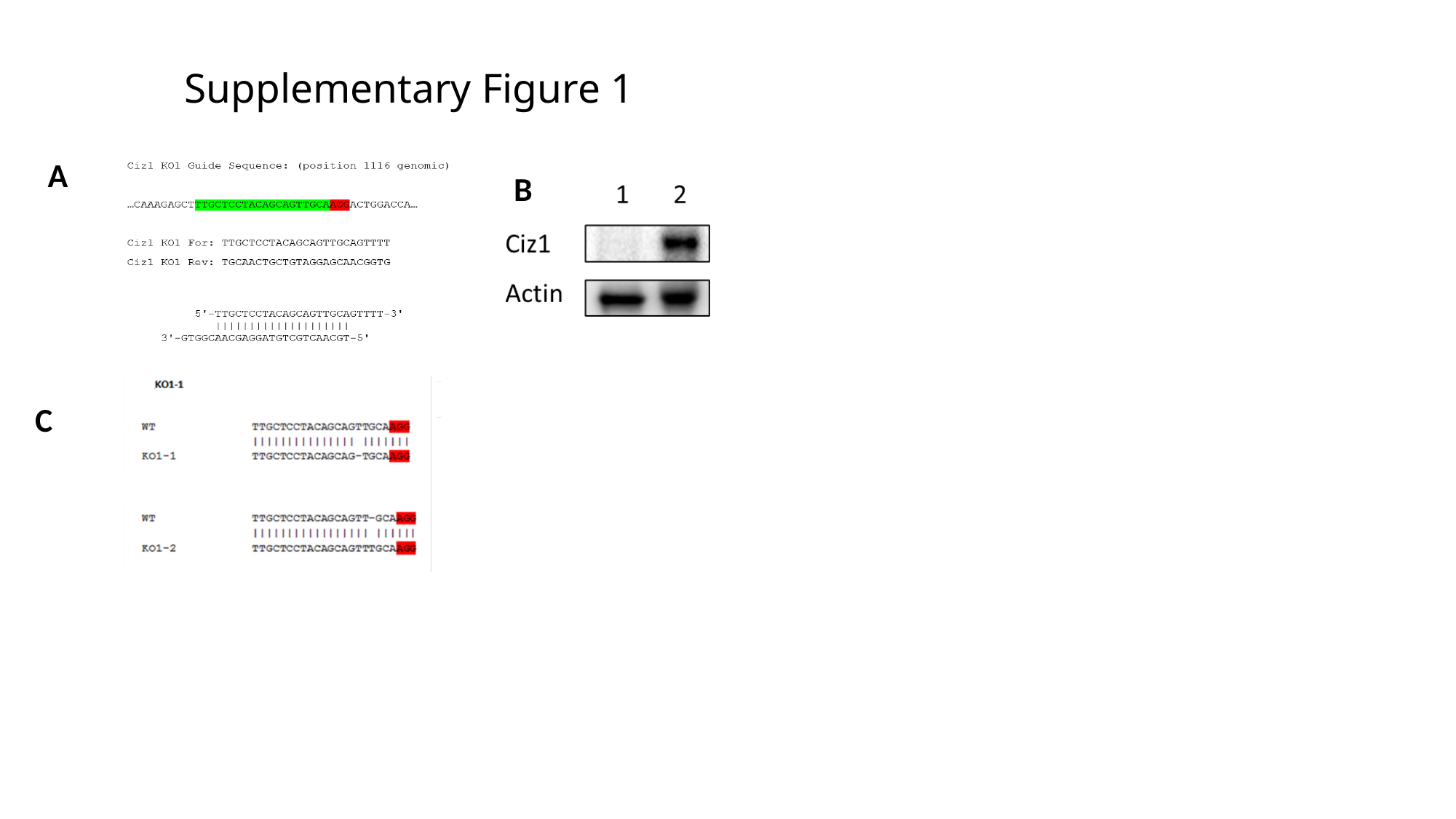

Supplementary Figure 1
#
A
B
C

### Figure S2

#### Slide 1
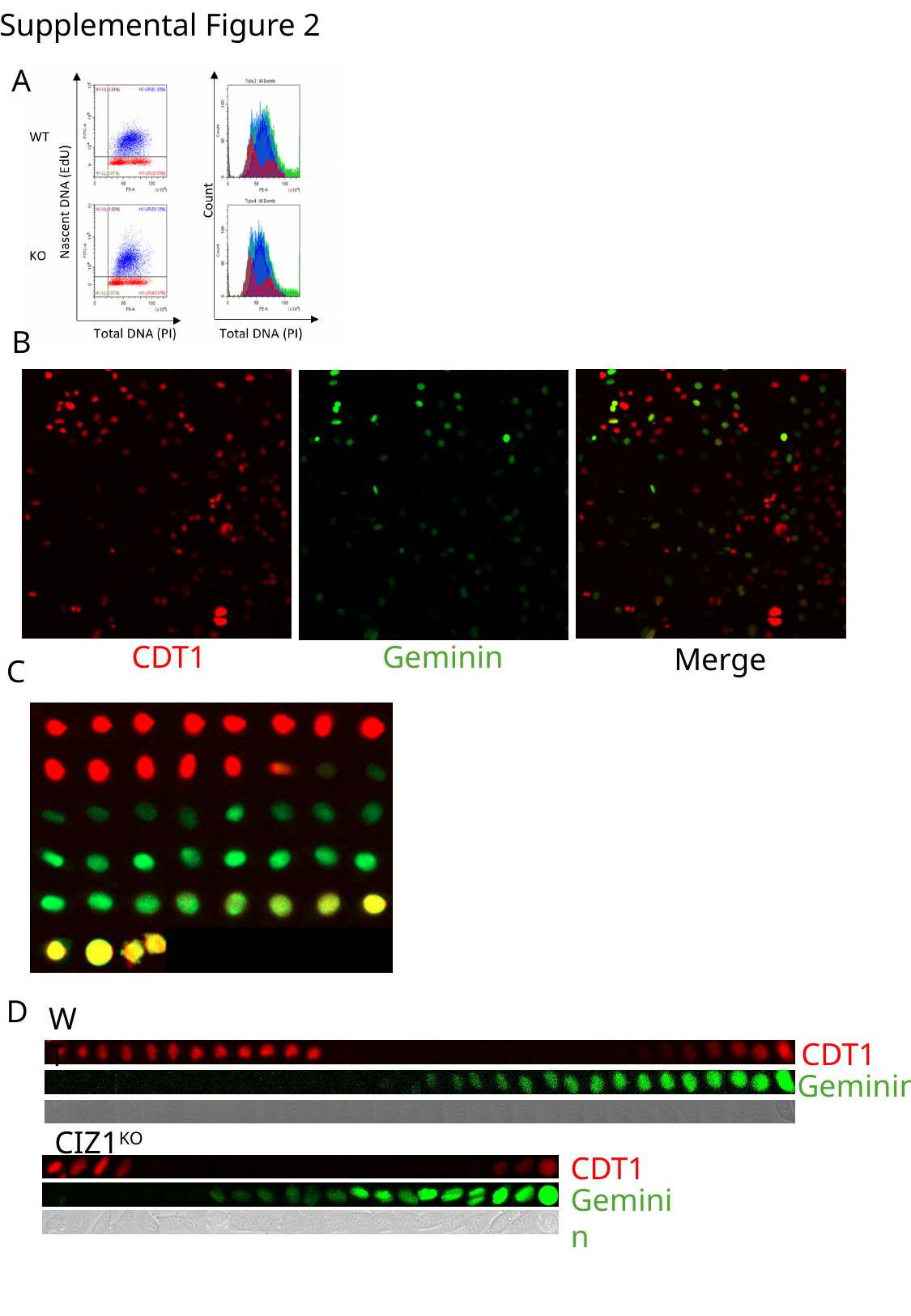

Supplemental Figure 2
A
B
Geminin
CDT1
Merge
C
D
WT
CDT1
Geminin
CIZ1^KO^
CDT1
Geminin

### Figure S3

#### Slide 1
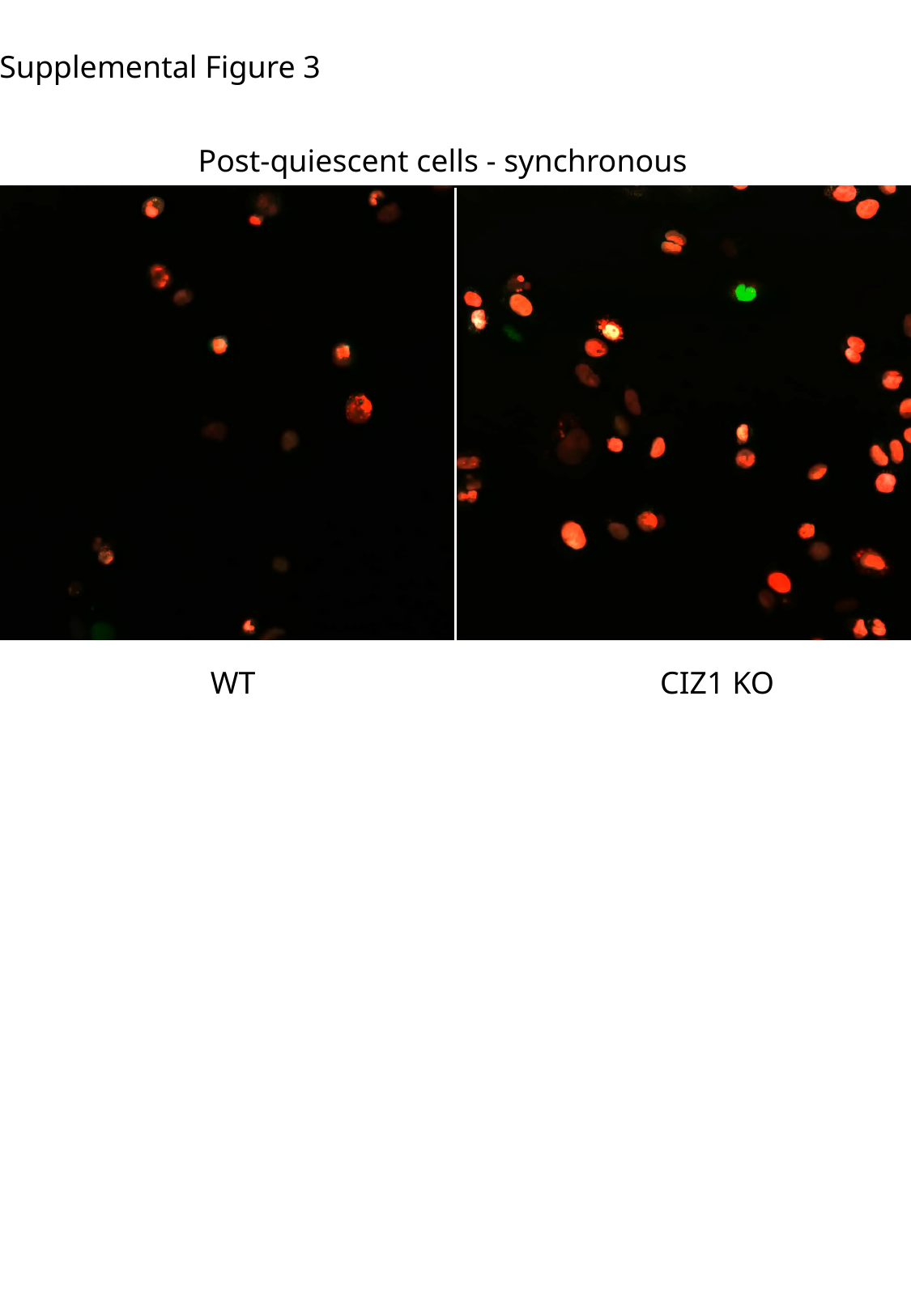

Supplemental Figure 3
Post-quiescent cells - synchronous
WT
CIZ1 KO
